# The fitness of *Pseudomonas aeruginosa* quorum sensing signal cheats is influenced by the diffusivity of the environment

**DOI:** 10.1101/082230

**Authors:** Anne Mund, Stephen P. Diggle, Freya Harrison

## Abstract

Experiments examining the social dynamics of bacterial quorum sensing (QS) have focused on mutants which do not respond to signals, and the role of QS-regulated exoproducts as public goods. The potential for QS signal molecules to themselves be social public goods has received much less attention. Here, we analyse how signal-deficient (*lasI*^−^) mutants of the opportunistic pathogen *Pseudomonas aeruginosa* interact with wild-type cells in an environment where QS is required for growth. We show that when growth requires a ‘private’ intracellular metabolic mechanism activated by the presence of QS signal, *lasI*^−^ mutants act as social cheats and outcompete signal-producing wild-type bacteria in mixed cultures, because they can use the signals produced by wild type cells. However, reducing the ability of signal molecules to diffuse through the growth medium, results in signal molecules becoming less accessible to mutants, leading to reduced cheating. Our results indicate that QS signal molecules can be considered as social public goods in a way that has been previously described for other exoproducts, but that spatial structuring of populations reduces exploitation by non-cooperative signal cheats.

**Importance:** Bacteria communicate via signaling molecules to regulate the expression of a whole range of genes. This process, termed quorum sensing (QS), moderates bacterial metabolism in many environmental conditions, from soil and water (where QS-regulated genes influence nutrient cycling) to animal hosts (where QS-regulated genes determine pathogen virulence). Understanding the ecology of QS could therefore yield vital clues as to how we might modify bacterial behaviour for environmental or clinical gains. Here, we demonstrate that QS signals act as shareable public goods. This means that their evolution, and therefore population-level responses to interference with QS, will be constrained by population structure. Further, we show that environmental structure (constraints on signal diffusion) alters the accessibility of QS signals and demonstrates that we need to consider population and environmental structure to help us further our understanding of QS signaling systems.

## Introduction

Bacterial quorum sensing (QS) is a cell-to-cell signalling mechanism that coordinates a range of behaviours at the population level (1, 2). QS facilitates density-dependent production of extracellular molecules including nutrient-scavenging enzymes and virulence factors. These molecules have been termed ‘public goods’ because their benefits can be shared by all cells in the local population (3–6). Because these QS-regulated exoproducts are metabolically costly for cells to produce, QS can also be exploited by non-cooperating “cheats”: cells that do not respond to QS signals and so pay no costs, but which exploit wild type populations because they benefit from the public goods produced by wild type neighbouring cells (4–7). Experiments into the social dynamics of QS have traditionally focused on these “signal blind” mutants, and a number of studies have shown that such mutants can arise in laboratory cultures and during infections (8–14). In various laboratory conditions and *in vivo* infection models, they have been shown to act as social cheats (6, 15–17).

However, little attention has been paid to whether QS signals themselves can act as exploitable public goods, despite there being a metabolic cost associated with the production of QS signals (18, 19). Here, we analyse how signal-negative (*lasI*-) mutants of the opportunistic pathogen *Pseudomonas aeruginosa* socially interact with wild-type cells in an environment where growth requires the cells to have a functional QS system, but where the fitness benefits of QS are ‘private’ to individual cells. *P. aeruginosa* regulates the production of many virulence factors through two *N*-acyl homoserine lactone (AHL) based QS systems. These systems, termed the *las* and *rhl* systems, produce and respond to the signals *N*-(3-oxododecanoyl)-L-homoserine lactone (3O-C12-HSL) and *N*-butanoyl-L-homoserine lactone (C4-HSL) respectively (1, 20). We conducted our experiments in a growth medium containing adenosine as a carbon source. Adenosine is deaminated to form inosine, which is degraded inside the cell by a nucleoside hydrolase (Nuh) to hypoxanthine plus ribose; hypoxanthine is then metabolised to produce glyoxylate plus urea (21). QS is crucial for growth in this medium because the *las* system (through the regulator LasR), positively regulates Nuh. Because Nuh acts intracellularly, any loss of fitness due to mutation of the signal gene *lasI* will be directly due to the lack of signal – not to any downstream effect on the production of extracellular enzymes. We demonstrate that, when provided with adenosine as a carbon source, *lasI*^−^ mutants act as cheats: they grow poorly in monoculture but have a higher relative fitness than the signal-producing wild type in mixed cultures. In contrast, *lasR* mutants, which cannot regulate Nuh in the presence of signal, do not gain any fitness benefits in mixed culture with wild type cells.

In contrast to experiments performed in well-mixed liquid medium in test tubes, interactions between bacterial cells in natural environments (including infections) are affected by spatial assortment and structuring (22–24). This affects how behaviours evolve (25–29). We tested how simple spatial structuring, through the addition of agar to the growth medium, alters QS signal diffusion and the social dynamics of wild-type and *lasI*^−^ cells. Consistent with work on other bacterial public goods (27), we demonstrate that the ability of lasI-mutants to cheat is significantly reduced in structured populations. These results have implications for understanding how and why bacterial signaling evolves, and the likely evolutionary fate of different types of QS mutant in varied environmental conditions (30).

## Materials and Methods

### Bacterial strains

The strains used were the wild type *Pseudomonas aeruginos*a laboratory strain PAO1 and isogenic mutants created via insertion of a gentamicin resistance gene in the QS genes *lasI* (PAO1 *lasI*::Gm; referred to as *lasI*^−^ (31)) or *lasR* (PAO1 *lasR*::Gm; referred to as *lasR*^−^ (16)). To test 3O-C12-HSL diffusion in different media, we used a reporter strain of the *lasI*^−^ mutant. This contains a chromosomal *luxABCDE* fusion to the promoter of the *lasB* gene, which encodes the QS-dependent protease LasB (PAO1 p*lasB::lux* (31)).

### Growth conditions

Quorum sensing medium (QSM) was modified from two previous studies (5, 32). QSM consisted of M9 Minimal Salts supplemented with 6.8 g/L Na_2_HPO_4_, 3 g/L KH_2_PO_4_, 0.5 g/L NaCl. 10µM NH_4_Cl, 0.1 µM CaCl_2_ and 1 µM MgSO_4_. QSM was supplemented with 0.1 % w/v of carbon sources as a mix of Casamino acids (CAA) and adenosine and the medium was filter sterilized. The exact ratio of CAA and adenosine was varied as detailed in the Results section. Liquid culture experiments were conducted in 24-well plates with a volume of 2 mL of media. Cultures were incubated overnight in LB medium at 37°C on an orbital shaker and standardised to an OD_600_ of 0.8–0.9; 2 µL of pure or mixed inoculum was added to each experimental culture. The starting frequency of the mutant was determined by diluting and plating the starter cultures to determine the number of colony-forming units (CFU) of each genotype. Experimental cultures in QSM were incubated at 37°C with orbital shaking for 24h or 48 h. After that time, cultures were diluted and replica plated on LB and LB + gentamycin (25 µg/ml) agar to enumerate the CFU of PAO1 and *lasI* or *lasR* mutants in mixed culture. Experiments in solid media were conducted in QSM + 2% w/v agar in 1 ml volumes in 48-well plates. Inoculation and culture conditions were otherwise identical to experiments in liquid medium. To break up agar prior to dilution and plating, the solid 1 ml agar cubes were retrieved from the plate and divided into thirds with a sterile metal spatula; each third was placed in a screw-cap tube containing 500 µl phosphate-buffered saline and 6 metal beads (Cambio) and homogenised using a FastPrep-24 5G bead beater (MP Biomedicals).

### Measures of signal concentration

To measure the concentration of QS signal (*N*-3-oxo-dodecanoyl-L-homoserine lactone; 3O-C12-HSL) present in 48 h cultures, 100 µl of each culture supernatant was mixed with 100 µl of a log phase culture of a luminescent *Escherichia coli* bioreporter (pSB1075; (33)) in the wells of a 96-well plate. This mixture was incubated for 4 h in a Tecan multimode plate reader and luminescence and OD_600_ recorded at 15-min intervals. To estimate 3O-C12-HSL concentration, the luminescence of experimental samples was compared against a calibration curve constructed using QSM supplemented with known concentrations of purified 3O-C12-HSL.

### Assaying the effect of agar on QS signal diffusion

Agar has been successfully used to retard the diffusion of other bacterial exoproducts (27). To verify that agar affects 3O-C12-HSL diffusion in QSM, and to determine the optimal agar concentration to use in further experiments, we devised a “sandwich experiment” in which a population of bacteria that switch on a luminescent reporter gene in response to QS signal, but which cannot themselves produce signal, were separated from a reservoir of purified signal by a layer of agar-supplemented medium. By measuring the time to expression of the luminescent reported, we can assess the extent to which the agar barrier delays diffusion of the signal from the reservoir to the reporter population. 0.1 ml LB supplemented with 0.5% w/v agar and containing 0.5 µM purified 3O-C12-HSL was added to the wells of a 48-well plate and allowed to solidify. A second layer of 0.8 ml LB supplemented with 1, 2, 3 or 4% w/v agar was then added on top of the signal-containing layer.

Each agar concentration was replicated in 6 wells. This layer was allowed to solidify and a final layer of LB containing 0.5% w/v agar and the reporter PAO1 *lasI*^−^ p*lasB*::*lux* (overnight culture at OD_600_ of 0.2) added. The plate was incubated in a Tecan multimode reader for 8 h and luminescence read at 10 min intervals. As shown in Figure S1, increasing agar concentration progressively delayed and reduced expression of luminescence. In order to check if higher luminescence was due to increased bacterial numbers, bacteria were retrieved and CFU were counted by plating. Median CFU was similar when 1 % or 2% agar was used (approx. 1.410^7^), but decreased by 30% when more agar was added (to approx. 110^7^). It was difficult to determine whether this was due to agar retarding growth at high concentrations, or simply due to the increased difficulty of thoroughly homogenising media rich in agar. 1% agar was therefore chosen for use in further experiments.

### Statistical analysis

Relative fitness of mutants, *v*, was calculated as *x*_2_(1-*x*_1_) / *x*_1_(1-*x*_2_), where *x*_1_ is the starting frequency of the mutant and *x*_2_ is the end frequency. It follows from the definition that a relative fitness <1 signifies a decrease in mutant frequency, while a relative fitness >1 signifies an increase in mutant frequency. Statistical analysis of the results was conducted in R 3.2.3 (34) using generalized linear models assuming an underlying gamma distribution, with adenosine treated as a continuous variable and block and treatment (liquid/solid medium) fitted as factors. Raw data for all analyses reported is supplied as Supplemental Material (Data S1).

## Results

### *las* mutants grow poorly in an environment where QS is required for growth

Previous work has shown that *las* mutants grow poorly in media where QS is required for growth (5, 7, 35). We first confirmed that both *lasI*^−^ and *lasR*^−^ mutants were reduced in fitness in the specific medium we chose for our experiments. We grew PAO1 and each mutant in a minimal medium base containing 0.1% w/v carbon source. The carbon source was composed of casamino acids (CAA, available for use by all cells, regardless of genotype) and adenosine (which can only be metabolised when QS is functional in cells), in varying ratios. As the relative amount of adenosine increased and cells were increasingly dependent on QS, the total cell density after 48 h was reduced, and this effect was more pronounced in *lasI* monocultures than in wild-type monocultures (Figure 1, CFU pure adenosine of CFU pure CAA: 4.6% for wild type, 0.1–0.2% for *las* mutants). When all the available carbon was supplied as adenosine, *lasI*^−^ monocultures grew to approximately 7% of the density of wild-type monocultures.

**Figure 1.**
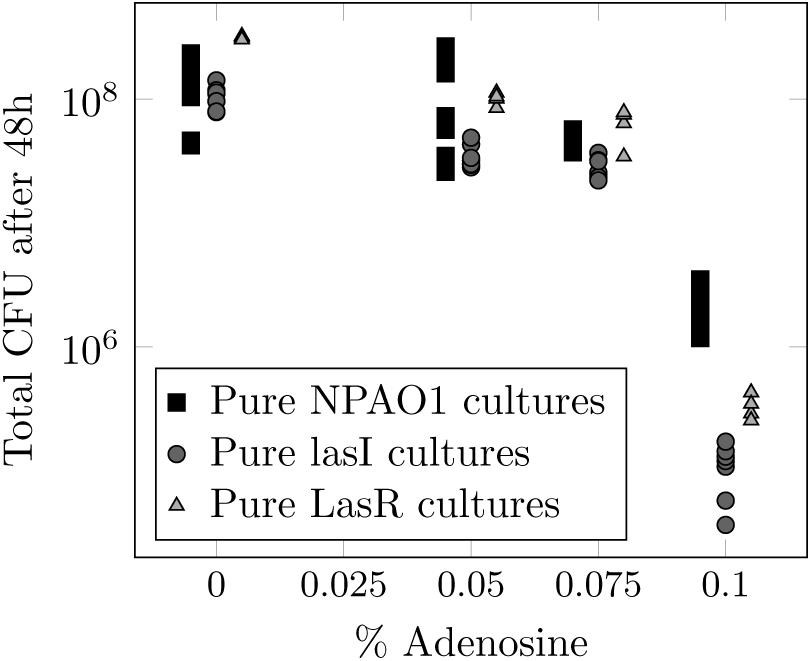
Population density (colony-forming units: CFU) after 48h of growth in quorum sensing medium with varying ratios of casamino acids:adenosine supplied to a total concentration of 0.1% w/v carbon source. Black dots: indvididual wild-type cultures; grey dots: indicate *lasI*^−^ cultures; white dots: *lasR* cultures.

### Signal-negative *lasI*^−^ mutants act as social cheats in adenosine-based growth medium, but signal-blind *lasR*^−^ mutant do not

We next tested whether adding purified 3O-C12-HSL, or co-culturing with signal-producing wild-type bacteria, could restore the growth of *lasI*^−^ mutants. We calculated the fitness of *lasI*^−^ mutants relative to the wild type (i) in pure culture with or without exogenous 3O-C12-HSL, and (ii) in 1:1 co-culture with wild-type PAO1. Experiments were conducted in quorum-sensing media with varying ratios of CAA and adenosine as above. A relative fitness of 1 signifies similar growth of mutant and wild type bacteria, while values <1 reflect relatively poorer growth of the mutant and values >1 reflect better growth of the mutant. Figure 2 shows fitted models describing how the relative fitness of *lasI*^−^ (a,b) and *lasR*^−^ (c,d) mutants are affected by culture conditions. The raw data to which the models were fitted are plotted in Figure S2.

**Figure 2.**
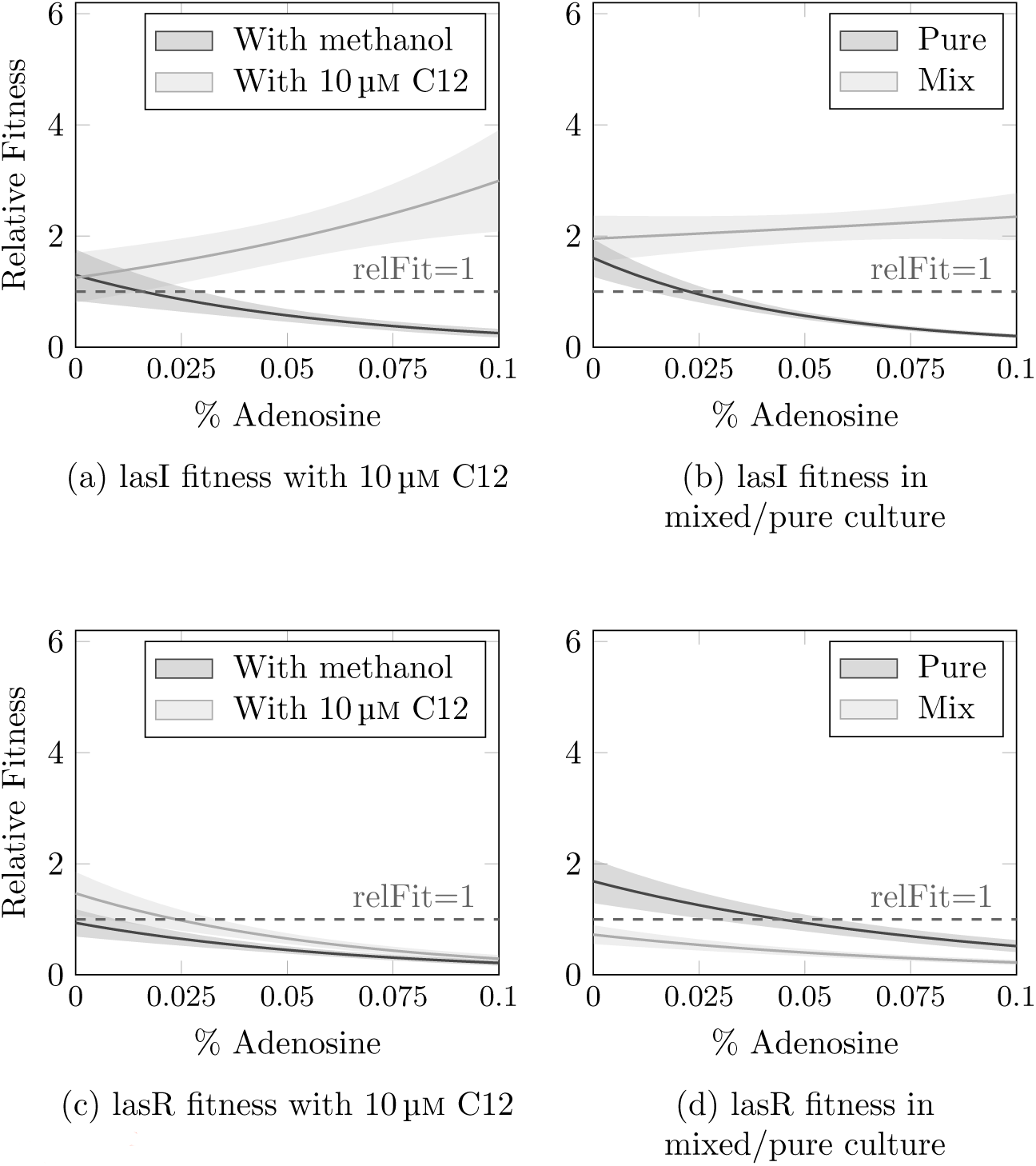
Results of fitting generalized linear models to relative fitness data from experiments with *lasI*^−^ (a,b) and *lasR*^−^ (c,d) mutants. Lines show fitted models, shaded areas denote standard deviation. Relative fitness is compared between pure cultures of each mutant with added 3O-C12-HSL or with a solvent-only control (a,c), and between pure cultures and mixed culture with wild-type bacteria (b,d). Raw data for these experiments are shown in Figure S2.

Pure *lasI*^−^ cultures became progressively less fit than the wild type as access to carbon depended more on quorum sensing (negative correlation between relative fitness and % adenosine: coefficient −21.6, *p* < 0.001). However, when 10 µM exogenous 3O-C12-HSL was supplied, *lasI*^−^ mutants surpassed the wild type in growth (positive correlation between relative fitness and % adenosine: coefficient 8,74, *p* = 0.003; Figure 2a, S2a) This result is consistent with previous work demonstrating a cost to 3O-C12 production (18). As predicted, this ability of *lasI*^−^ mutants to use exogenous signal, combined with the cost of signal production to the wild type, means that *lasI*^−^ mutants grown in co-culture with wild type bacteria act as social cheats: the average relative fitness was consistently > 1 and did not decline as % adenosine increased (coefficient 1.8, *p* = 0.88 Figure 2b, S2a). There was, however, a slight drop in relative fitness when all carbon was supplied as adenosine (Figure S2a. This is most likely attributable to the wild-type bacteria growing more slowly and taking longer to fully switch on QS responses. Both wild-type growth (Figure 1) and the pool of available signal (Figure S3) are reduced in this condition, leaving less opportunity for exploitation by cheats.

Post-hoc comparisons confirmed that *lasI*^−^ relative fitness was significantly increased in mixed populations vs. pure cultures in all media containing adenosine (*t*-tests, *p* < 0.01). When all carbon was supplied as CAA and signal is not required, there was no significant effect of pure vs. mixed culture on fitness (*p* > 0.95) Taken together with the fact that mixed cultures grew to a lower density than the wild-type cultures (Figure S5), these results indicate that *lasI* mutants have an increased fitness when grown in the presence of wild-type bacteria under conditions requiring social interaction, while in turn decreasing wild-type fitness.

To ensure that the effect described above was due to the social dynamics of signal production, and not to the well-documented social dynamics of downstream exoproducts, this experiment was repeated using a *lasR*^−^ mutant. *lasR*^−^ mutants are unable to respond to 3O-C12-HSL and should therefore not be able to derive fitness benefits from exogenous signal in our setup. We found the same negative correlations between % adenosine and CFU (Figure 1) and % adenosine and fitness relative to the wild type (Figure 2c, S2b, coefficient −12.8, *p* = 0.001) as with the *lasI*^−^ mutant for *lasR*^−^ monocultures. Crucially, *lasR*^−^ relative fitness was not rescued by adding 3O-C12-HSL or by co-culturing with the wild type (coefficient −10.7, *p* < 0.001; Figure 2c,d and S2b): i.e. these mutants could not exploit wild type bacteria.

### Slowing signal diffusion reduces *lasI*^−^ mutant cheating

As a last step, we tested whether impeding the diffusion of signal molecules would make the *lasI*^−^ mutant less effective as a cheat. Reduced diffusion was achieved by adding agar to solidify the growth medium (Figure S1). *lasI*^−^ monocultures showed comparable declines in fitness in liquid and solid medium (Figure 3a, S4a. ANOVA: liquid/solid F_(1,124)_ = 6.13, *p* = 0.01; adenosine F_(1,123)_ = 117.58, *p* < 0.001; interaction F_(1,122)_ = 0.1658, *p* = 0.68). In mixed cultures, the relative fitness of the *lasI*^−^ mutant was positively correlated with the percentage of carbon available as adenosine in liquid cultures, as expected under cheating: but when the medium was solidified, *lasI*^−^ relative fitness actually showed a modest negative correlation with percentage adenosine (Figure 3b, S4b. ANOVA: liquid/solid F_(1,123)_ = 11.1324, *p* = 0.001; adenosine F_(1,124)_ = 0.0236, *p* = 0.88; interaction F_(1,122)_ = 0.9760, *p* = 0.33). This demonstrates that there is less opportunity for cheating in an environment where signal molecules cannot diffuse freely.

**Figure 3.**
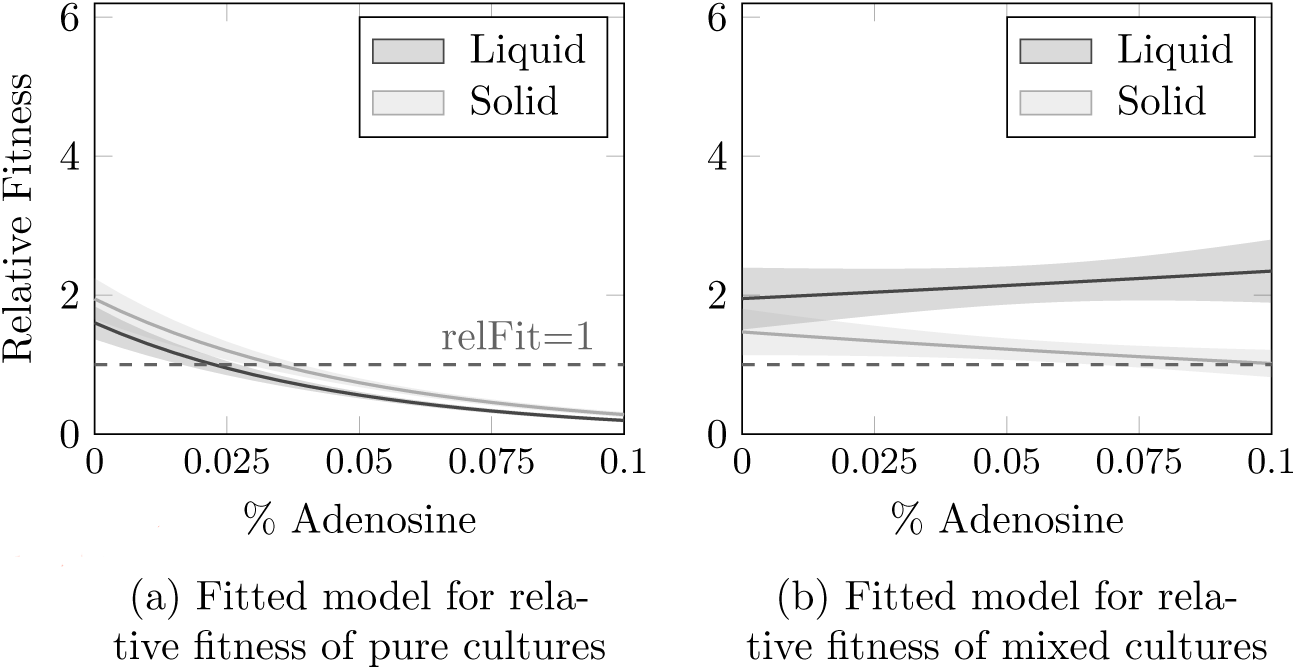
Comparison of *lasI*^−^ relative fitness in pure culture (a) or mixed culture with the wild type (b). Results of fitting generalized linear models to relative fitness data from experiments in liquid *versus* agar-supplemented medium. Lines show fitted models, shaded areas denote standard deviation. Raw data for these experiments are shown in Figure S4.

## Discussion

While there has been some discussion of QS signals as public goods (e.g. (36)), most published work on the social evolution of QS focuses on signal-blind mutants and the benefits of cheating on the production of QS-regulated exoproducts(5, 7, 14, 32). Here we provide the first direct evidence that, in addition to regulating the production of public goods, QS signal molecules are themselves capable of acting as public goods. Social cheating in the context of QS can therefore take multiple forms, depending on the environmental circumstances in which bacteria find themselves. Previous research had shown that (a) signal-blind mutants can be cheats when growth depends on the production of QS-dependent extracellular enzymes; and (b) cheating by signal-blind mutants can be constrained when some ‘private’ QS-controlled processes contribute to growth (35). Following recent confirmation that QS signals are costly to make (18), we now add two new perspectives to the social evolution of QS: (c) that signal-negative mutants can be cheats when growth depends entirely on private goods, regardless of any downstream effects on exoenzyme production; and (d) that this signal cheating can only occur when the environment permits sufficient diffusion of signal molecules.

A *lasI*^−^ mutant grew poorly in monoculture, but growth could be rescued by adding either purified 3O-C12-HSL signal or by co-culturing with wild-type, signal-producing bacteria. In co-cultures, the fitness of *lasI*^−^ mutants relative to the wild type increased as we forced the bacteria to rely more on adenosine for carbon. As the population became more reliant on QS, *lasI* mutants gained a greater fitness pay-off from exploiting costly, diffusible signal produced by the wild-type. *lasR*^−^ mutants did not gain a similar advantage from co-culture with the wild type in adenosine medium. These signal-blind mutants can take up signals, but cannot respond to them and so cannot switch on expression of the *nuh* hydrolase required for growth on adenosine (21, 35). The contrasting results for the two different QS mutants confirm that, in this environment, the QS signal itself acts as a public good (Figure 4)

**Figure 4.**
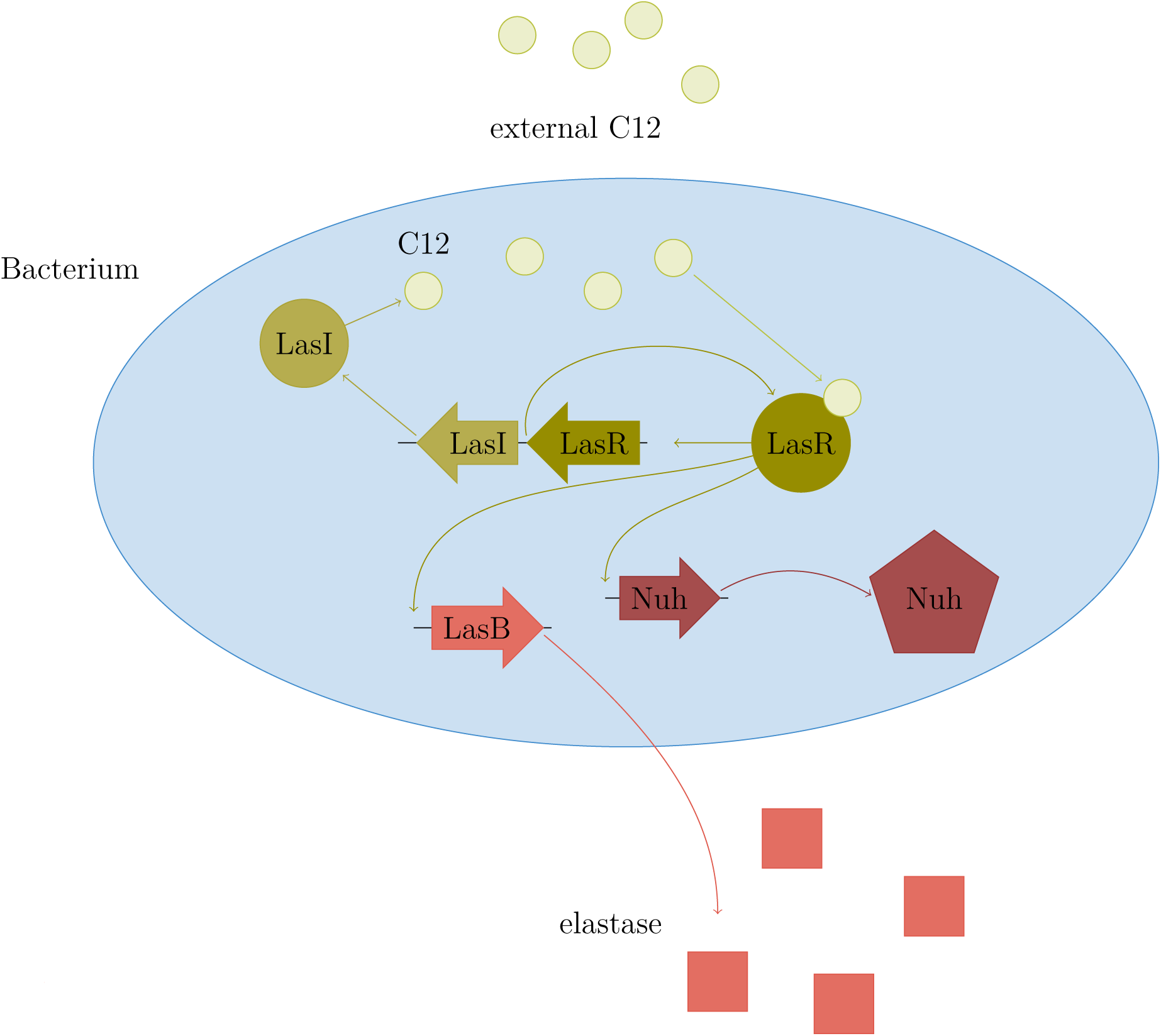
Schematic of the effects of LasIR quorum sensing on the production of intracellular and extracellular enzymes by *Pseudomonas aeruginosa*.

Adding agar to the growth medium lowered both the diffusion of signal molecules and the relative fitness of cheats in mixed culture. Thus adding simple spatial structuring into our system had a significant impact on the ability of signal-negative mutants to cheat on the wild type. There was no effect of structuring on relative fitness in pure cultures: even though agar enhanced the overall growth rate of the bacteria, the basic costs and benefits of signalling remained the same in liquid and solid media. This final observation is consistent with work on other bacterial public goods (27). We thus predict that the evolution of QS signalling strategies will be influenced by population genetic and spatial structure, and that signal-negative cheats might only rise to appreciable frequencies in environments where signals diffuse freely. For example, the thick, adhesive mucus and bacterial biofilm polymers that block the airways of cystic fibrosis patients with chronic lung infection may partially protect producers from cheating by signal-negative mutants (24). Spatial dynamics play a huge role in the real-life ecology of environmentally and clinically important microbial ecosystems, and are therefore of considerable interest to microbiologists investigating the roles of bacteria in processes as diverse as geochemical cycling, soil health, fouling and infection (29, 37, 38).

Our work opens up new avenues for exploring how, when and why bacterial signalling evolves in different environments and why we find a variety of QS mutant genotypes and phenotypes in natural environments (8–14, 39, 40). Given what we are now learning about the evolution of traits such as QS and how spatial structure changes the evolutionary dynamics, we suggest that there is a need to carefully consider the experimental design of *in vitro* experiments to increase their relevance to actual infections (41). To be forewarned is to be forearmed: a more accurate understanding of microbial ecology and evolution, gained from more realistic lab experiments, will be a vital weapon in the fight against antibiotic resistant infection.

## Acknowledgements

AM was sponsored by the Erasmus+ program. This work was supported by a Human Frontier Science Program Young Investigators grant to SPD (RGY0081/2012), a NERC grant to SPD (NE/J007064/1), as well as by the German Research Foundation (DFG) and the Technical University of Munich (TUM) in the framework of the Open Access Publishing Program. We thank Alex Truman for supplying purified 3O-C12-HSL; Sheyda Azimi and James Gurney for helpful comments on the experimental work; and Burkhard Hense for useful discussion.

## Figure legends

**Figure S1.**
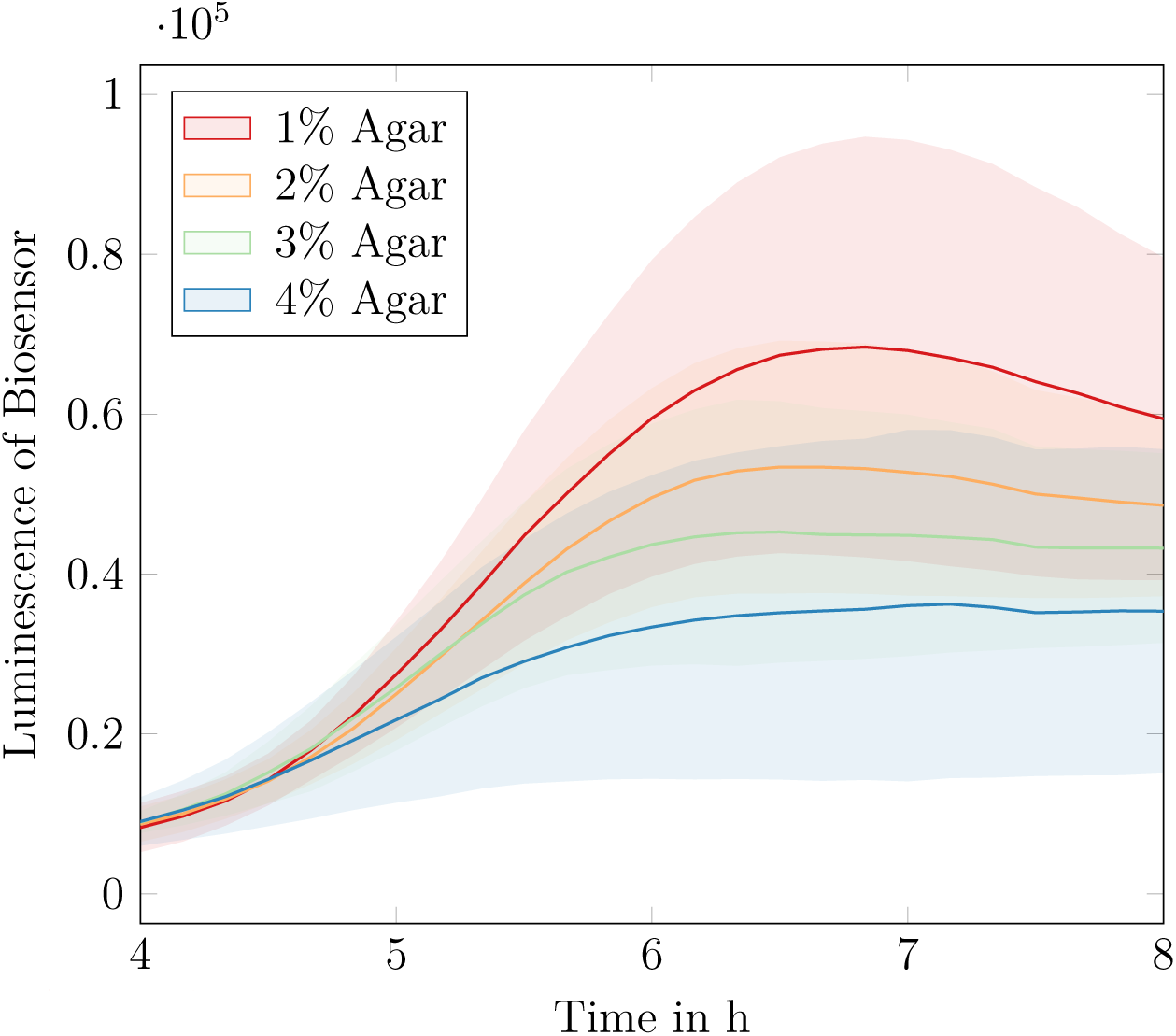
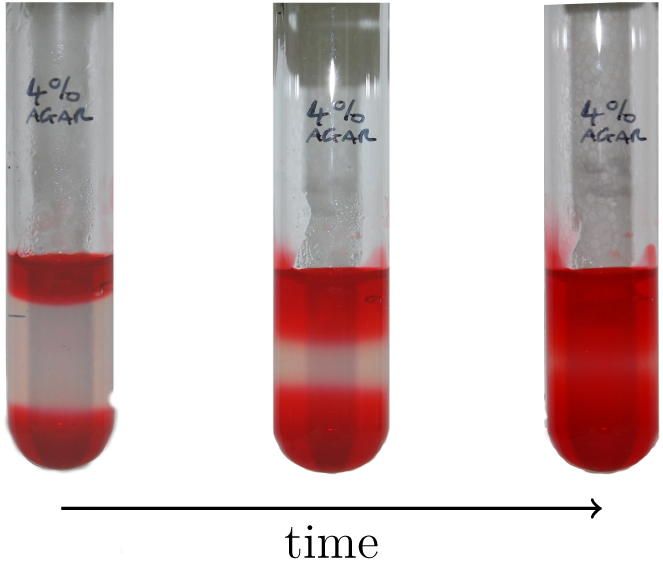
Supplementing growth medium with agar retards diffusion of 3O-C12-HSL. (a) Expression of a 3O-C12-HSL-dependent luminescent reporter construct is delayed and reduced when signal-negative reporter bacteria are separated from a reservoir or purified signal by an agar barrier. Lines and shading show means and s.d. of 15 replicates. (b) The experimental set-up, as described in the Materials and Methods, is easily visualised by replicating the experiment on a larger scale and adding red food colouring in place of bacterial signal.

**Figure S2.**
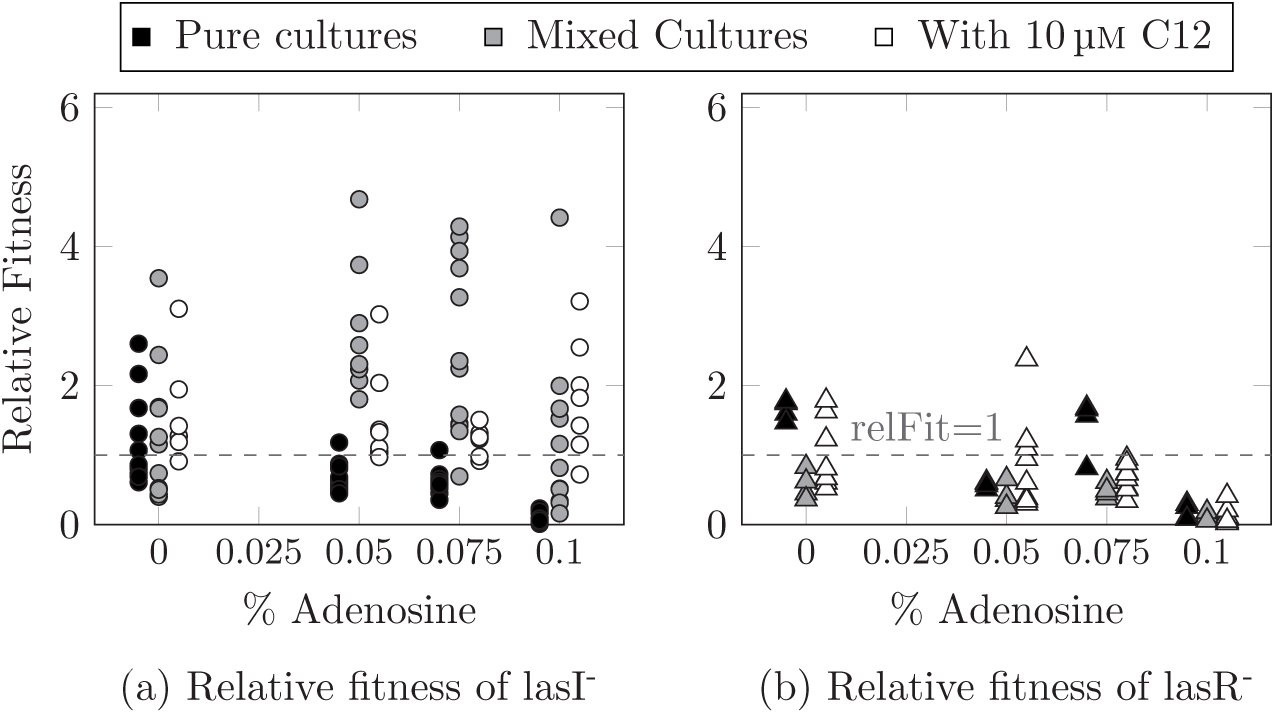
Relative fitness of (a) *lasI*^−^ mutants and (b) *lasR*^−^ mutants when in pure culture (black circles), grown in 1:1 mixture with wild type (grey circles) and in pure culture supplemented with 10 µM 3O-C12-HSL (white circles).

**Figure S3.**
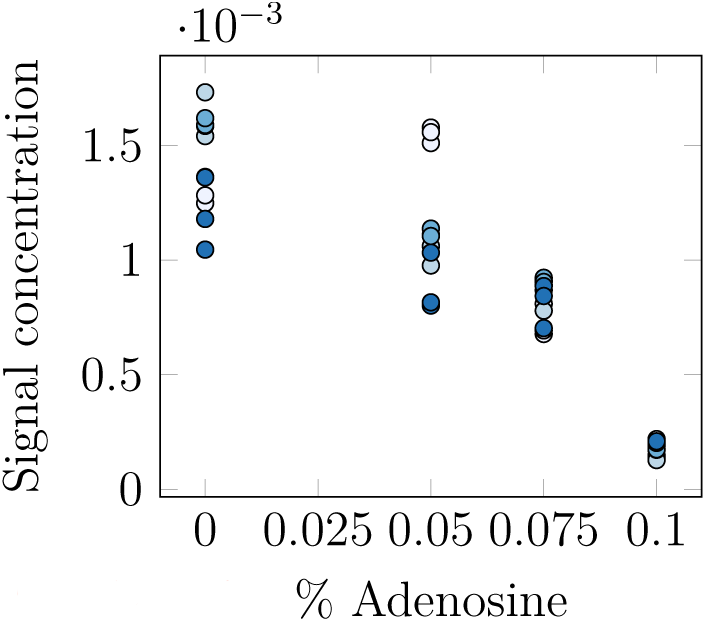
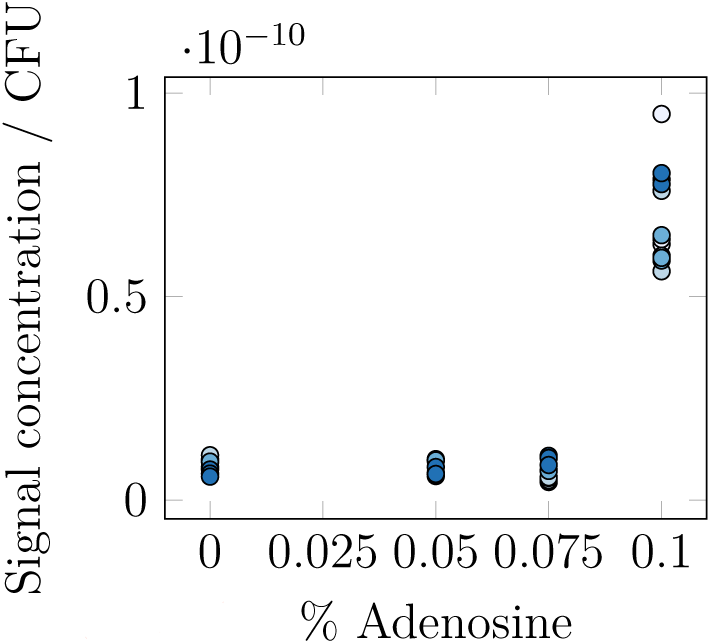
Measurements of 3O-C12-HSL concentration in wild-type supernatant after 48h. (a) The total concentration of 3O-C12-HSL declines with increasing adenosine. (b) After dividing the signal concentration by the CFU of the producing culture, one observes the reverse trend – concentration of 3O-C12-HSL per CFU increases with adenosine level and reaches its maximum in pure adenosine.

**Figure S4.**
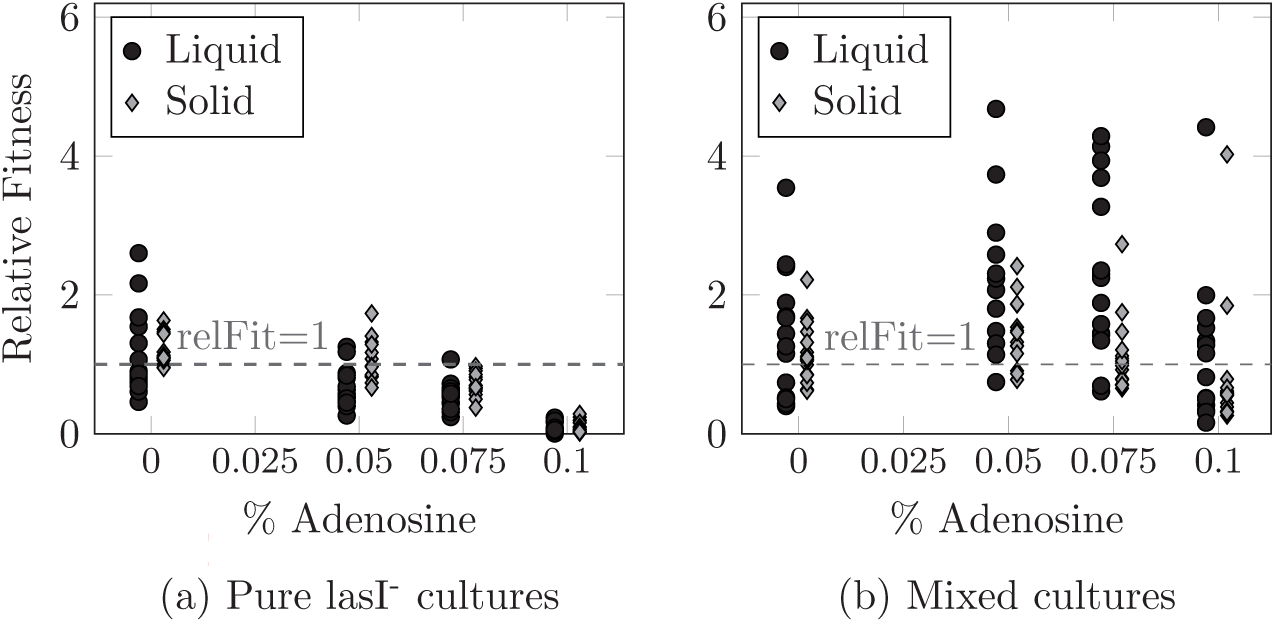
Relative fitness of *lasI*^−^ mutants in liquid medium (black circles) *versus* agar-supplemented medium (grey circles), (a) in pure culture and (b) in mixed culture with the wild type.

**Figure S5.**
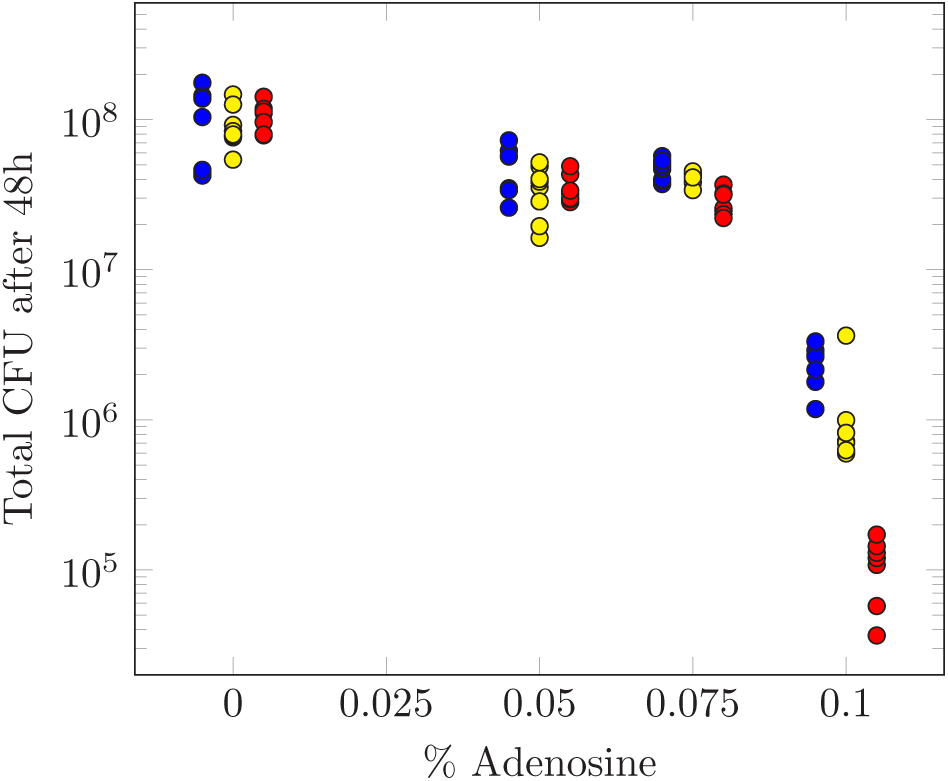
Population density (colony-forming units: CFU) after 48h of growth in quorum sensing medium, Blue dots: indvididual wild-type cultures; yellow dots: indicate *lasI* and wild-type mixed cultures; red dots: *lasI* cultures.

**Data S1.** Data for all analyses reported in the text will be made available on with the published version of this article.

